# Shaping neonatal immunization by tuning delivery of synergistic adjuvants via nanocarriers

**DOI:** 10.1101/2022.06.03.494753

**Authors:** Soumik Barman, Francesco Borriello, Byron Brook, Carlo Pietrasanta, Maria De Leon, Cali Sweitzer, Manisha Menon, Simon D. van Haren, Dheeraj Soni, Yoshine Saito, Etsuro Nanishi, Sijia Yi, Sharan Bobbala, Ofer Levy, Evan A. Scott, David J. Dowling

## Abstract

Adjuvanted nanocarrier-based vaccines hold substantial potential for applications in novel early-life immunization strategies. Here, via mouse and human age-specific *in vitro* modelling, we identified the combination of a small molecule STING agonist (2′3′-cyclic GMP-AMP, cGAMP) and a TLR7/8 agonist (CL075) to drive synergistic activation of neonatal dendritic cells and precision CD4 Th cell expansion via the IL-12/IFNγ axis. We further demonstrate that vaccination of neonatal mice with quadrivalent influenza recombinant hemagglutinin (rHA) and an admixture of two polymersome (PS) nanocarriers separately encapsulating-cGAMP (cGAMP-PS) and CL075 (CL075-PS) drove robust T helper (Th)1 bias, high frequency of T follicular helper (T_FH_) cells and germinal center B (GC B) cells along with IgG2c-skewed humoral response *in vivo*. Dual loaded cGAMP/CL075-PSs did not outperform admixed cGAMP-PSs and CL075-PSs *in vivo*. These data validate an optimally designed adjuvantation system, via age-selected small molecule synergy and a multi-component nanocarrier formulation, as an effective approach to induce type 1 immune responses in early life.

## INTRODUCTION

New vaccine strategies and formulations are needed to provide improved protection for vulnerable populations, such as infants and older adults (i.e., elderly)^1^. Adjuvants are critical components of vaccine formulations that can significantly increase immunization efficacy, primarily by functioning as agonists of pattern-recognition receptors (PRR) to stimulate innate immune cells. However, most adjuvanted vaccine formulations are not rationally selected or tailored to account for age-associated differences in innate immune responses, often resulting in suboptimal hyporesponsiveness and/or immunosenescent differences in translational vaccine outcomes^2, 3^. Strategies under consideration to address these distinct responses include a) combining two or more PRR agonists into a single vaccine,^4^ and b) optimization of immune-engineering strategy and delivery system design in an age-specific manner^5, 6^. Immunomodulation and intracellular delivery of antigen/adjuvant by nanocarriers comprised of poly(ethylene glycol)-*b*-poly(propylene sulfide) (PEG-*b*-PPS) block copolymers have been well-characterized^7–11^. PEG-*b*-PPS nanocarrier platforms not only permit efficient loading of both hydrophobic and hydrophilic adjuvants^7–11^ but also prime T cells by enhancing their uptake and presentation by antigen presenting cells (APCs)^7–11^.

Our group has previously employed adjuvant-loaded PEG-*b*-PPS polymersome (PS) nanocarriers^12^ to enable delivery to specific subsets of leukocytes (i.e., directly to APCs) and even specific subcellular compartments (i.e., endosomes). This strategy is highly advantageous for selective targeting of endosomal receptors, such as Toll-like receptor (TLR)7 and TLR8, and cytosolic receptors like stimulator of interferon genes (STING)^13^. We demonstrated that TLR7/8 agonist PS formulations mimic the immunomodulating effects of the live attenuated vaccine *Bacille Calmette-Guèrin* (BCG) which is commonly given to newborns in tuberculosis-endemic countries, by enhancing both innate and adaptive immune responses^12^. Notably, when co-loaded with the *Mycobacterium tuberculosis* antigen 85B peptide 25, the TLR8-agonist loaded PSs were comparable to BCG in inducing antigen-specific immune responses in hTLR8-expressing humanized neonatal mice *in vivo*^12^. Furthermore, we discovered that cGAMP is a potent activator of newborn DCs^13^. As compared to alum or cGAMP alone, immunization with cGAMP adsorbed onto alum via inherent phosphonate groups, enhanced murine newborn rHA-specific IgG2a/c titers, an antibody (Ab) subclass associated with the development of interferon (IFN)γ-driven type 1 immunity *in vivo* and endowed with higher effector functions^13^. Hence, we hypothesized that targeting both STING and TLR7/8 pathways in early life innate immune cells could be a promising precision vaccinology strategy.

In the present study, we identify that a combination of small molecules cGAMP and CL075 drive synergistic and robust activation of neonatal dendritic cells and precision CD4 Th cell expansion, via the IL-12/IFNγ axis *in vitro*. Then, we took advantage of the standardized PS platform to formulate separate cGAMP and CL075 loaded PSs (cGAMP-PS and CL075-PS) and studied tunable aspects of immunomodulation in early life immunization with the help of the quadrivalent influenza subunit vaccine, Flublok *in vivo*. We also co-encapsulated CL075 and cGAMP inside PSs and assessed differences in the resulting kinetics and immunostimulatory effects. Admixture of CL075-PS and cGAMP-PS primed rHA specific T_FH_ and GC B cell responses and induced robust humoral and rHA-specific Th1 and CD8^+^ T cell responses. By using the PS platform and the combination of STING and TLR7/8 agonists, we successfully enhanced rHA specific type 1 immunity in infant mice. Within a framework of precision vaccinology, formulation of such precision adjuvant delivery systems to fine tune vaccine immunogenicity may inform the development of next generation rationally designed vaccines for vulnerable pediatric and elder human populations.

## RESULTS AND DISCUSSION

### Dual STING and TLR7/8 stimulation drives IL-12 dependent Th1-Polarization in early life

STING^13–15^ and TLR7/8^12, 16, 17^ agonists induce overlapping but distinct innate immunological profiles^18^, with both upstream activation pathways leading to the secretion of innate type 1 interferon and pro-inflammatory cytokine production. These cytokines are critical to APC priming of T cells and the initiation of adaptive immune responses. To assess the immunostimulatory profiles and synergistic effects of cGAMP and CL075 on primary dendritic cells (DCs) from murine and human origins, we stimulated adult and neonatal bone marrow-derived dendritic cells (BMDCs) from mice (Figure 1A) and human monocyte derived dendritic cells (MoDCs) (Figure 1B) with increasing concentrations of each small molecule agonist for 24 h. We focused on TNF, IFNβ and IL-12p70 as surrogates for the capacity to induce a Th1-promoting innate response. CL075 and cGAMP synergistically induced concentration-dependent TNF and IFNβ production in neonatal and adult BMDCs (Figure 1A). A similar pattern was observed for the production of TNF and IL-12p70 by neonatal and adult human MoDCs (Figure 1B). Overall, the highest degree of synergy was noted for IL-12p70. Specifically, combined treatment with cGAMP and CL075 overcame neonatal DC hyporesponsiveness for IL-12p70 production, of which neonatal DC are known to have a limited production ability in response to most PRR agonists^19–21^. This hyporesponsiveness, combined with a baseline Th2-polarized response^22, 23^ ^24^, contributes to the distinctness of neonatal DCs.

**Figure 1.**
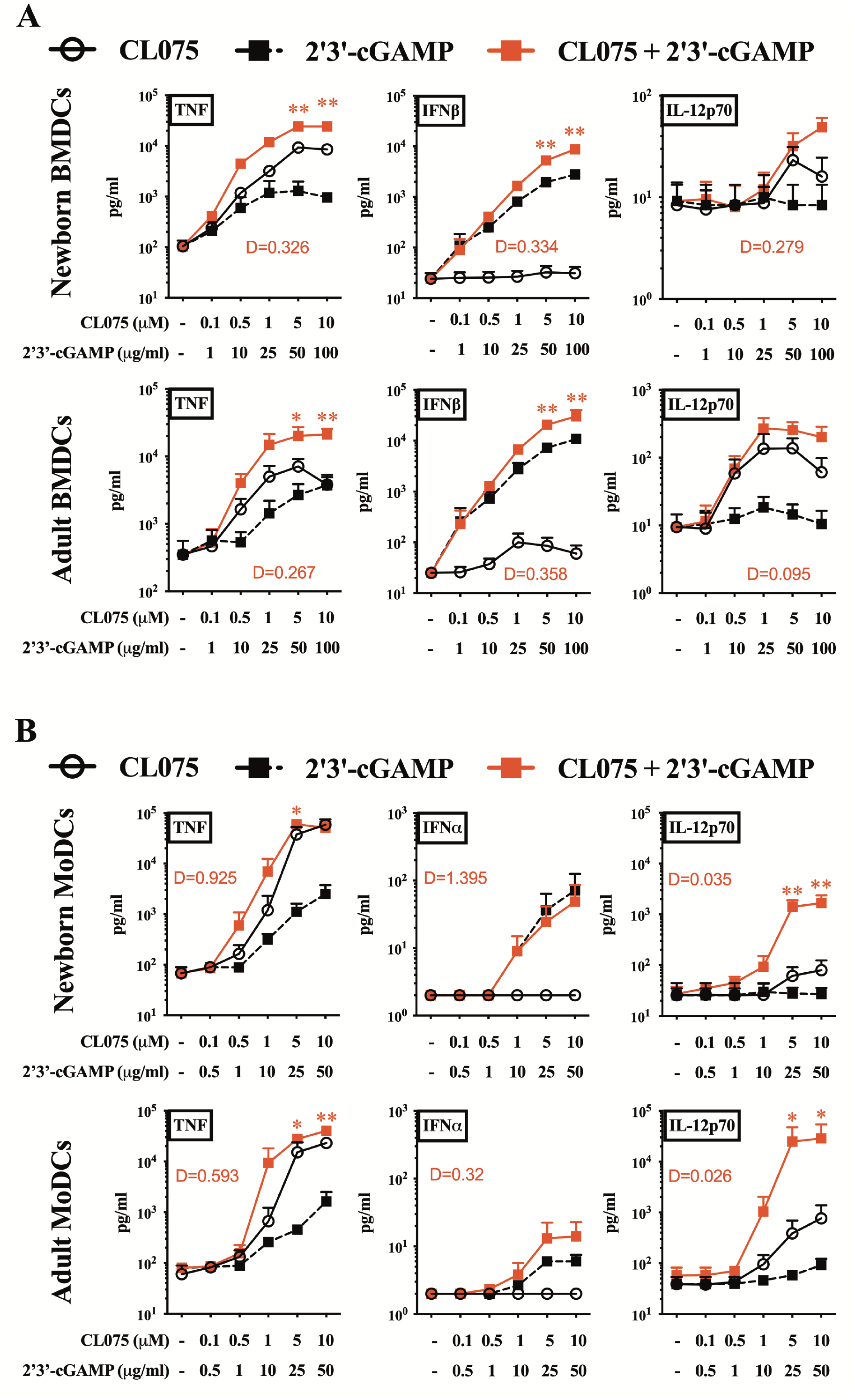
cGAMP and CL075 synergistically induced Th1 related cytokines in murine and human dendritic cells. (A) BMDCs from newborn (7 days of life) and adult (∼8 weeks old) mice and (B) MoDCs from human neonatal CBMC and adult PBMC were stimulated with the indicated concentration of 2’3’-cGAMP and CL075. Degree of synergy (1/D) was compared between groups. Data are representative of three independent experiments. Statistical comparison employed test one-way ANOVA; \**p* < 0.033, \*\**p* < 0.002, \*\*\**p* < 0.001 (n = 7 per group).

Next, we employed a 96 h human newborn T-helper polarization assay, which leverages the intrinsic characteristics of the newborn T cell compartment (composed mainly of naïve T cells) to iteratively characterize how adjuvant and nanocarrier formulation modulate T cell polarization in mixed mononuclear cell culture in the presence of a TCR-mediated stimulus. Cord blood mononuclear cells (CBMCs) were stimulated with αCD3 (polyclonal T cell activator) ± CL075 (5 μM), cGAMP (25 μg/mL) or combined, and the selective contribution of different factors (e.g., type I IFNs, IL-12p70) to T cell polarization was investigated via blocking Abs. Interestingly, and in line with DC data, we observed a restored Th1 predominance in neonatal leukocytes via the combination of CL075 and cGAMP (Figure 2A). To investigate further, we confirmed the accumulation of IFNγ^+^ CD4^+^ (Th1-polarized) neonatal T cells, minimal IL-4^+^ but not IL-17^+^, in the presence of both CL075 and cGAMP by flow cytometry (Figure 2B-E). The IFNγ response was significant both without or with TCR stimulation. IFNγ^+^ CD8^+^ neonatal T cells were detected upon combined stimulation of TLR7/8 and STING agonists (Figure 2B, F-H), with significant changes observed only in the presence of TCR stimulation. Importantly, using neutralizing Abs (nAbs) against Type I IFN signaling (αIFNAR2) and combined nAbs against IL-12p40/70 (αIL-12p40/70), we demonstrated that secretion of IFNγ^+^ by CBMCs over 96 h is reliant on innate IL-12 signaling (Figure 2I-J).

**Figure 2.**
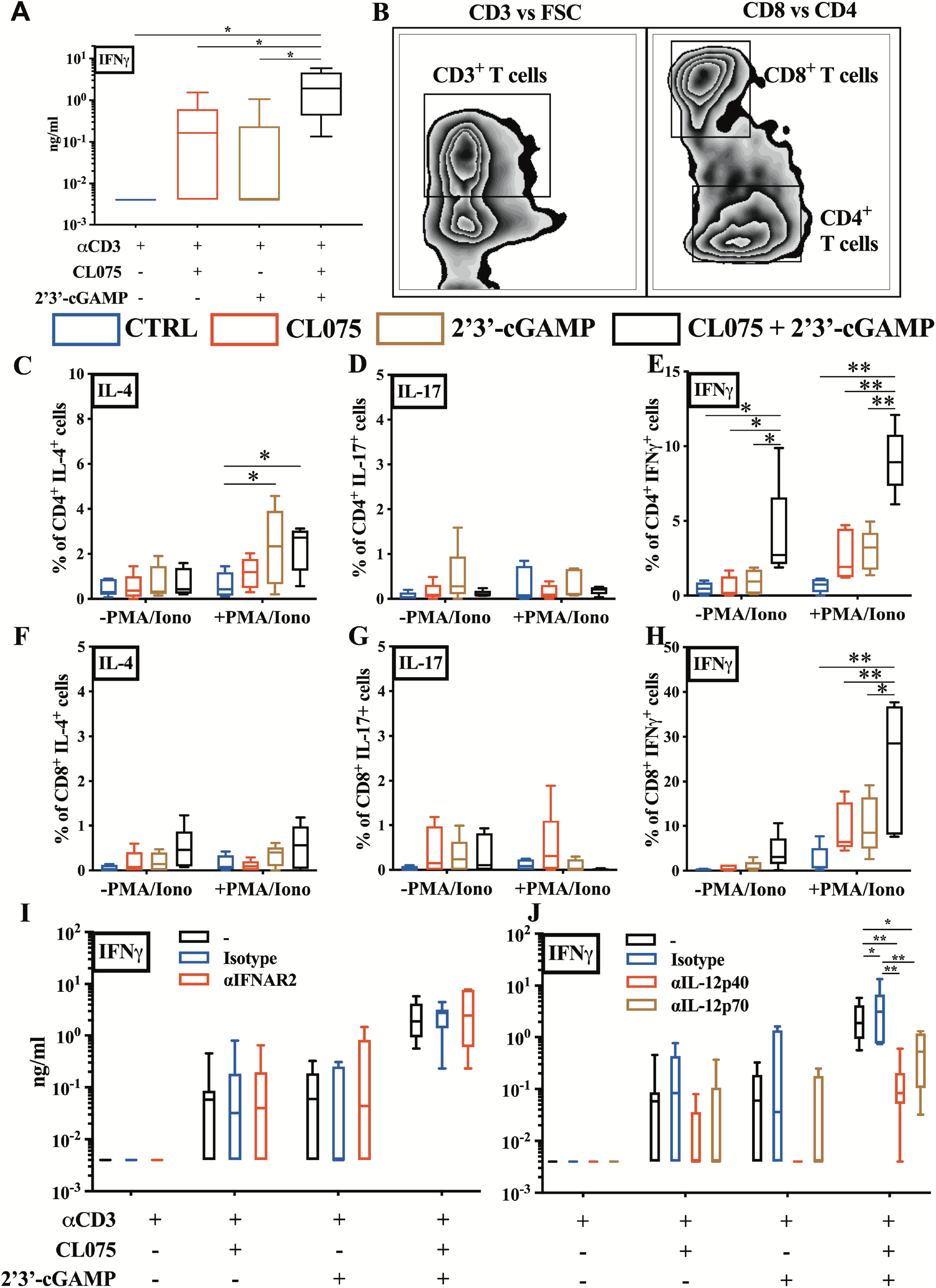
A combination of cGAMP and CL075 drove IL-12 dependent Th1 polarization in human neonatal T cells. (A) Neonatal CBMCs were cultured *in vitro* for 96 h in the presence of polyclonal T cell activator ℻-CD3 (1 μg/mL) with or without cGAMP (5 μM), CL075 (25 μg/mL) or cGAMP+CL075, followed by IFNγ production evaluation by ELISA. (B) Example gating strategy for quantification of CD4^+^ and CD8^+^ T cells (C-H) CBMCs were cultured *in vitro* as in A, but with the addition of cytokine blocker for the last 6h. After stimulation, cells were harvested, stained (intracellular cytokine staining) and analyzed by flow cytometry to quantify the percentage of T cells producing IL-4, IL-17 and IFNγ. (I) CBMCs were cultured *in vitro* as in A, but with the addition of human blocking Abs, ℻IFNAR2 or (J) ℻IL12p40/70. After stimulation, collected supernatant was evaluated via ELISA for IFNγ. Data were representative of two independent experiments. Statistical comparison employed test one-way ANOVA; \**p* < 0.033, \*\**p* < 0.002, \*\*\**p* < 0.001 (n = 7 per group).

### Synthesis and characterization of adjuvant-loaded PS nanocarriers

PEG-*b*-PPS copolymers with hydrophilic block weight fractions of 0.38 were self-assembled into PS as previously described^25, 26^. PSs were formed and loaded using the flash nanoprecipitation technique^7, 25^, with the hydrophobic CL075 loaded within the PS bilayer membrane and hydrophilic cGAMP encapsulated within the aqueous core (Figure 3A). Each batch of PSs was well-characterized, including quantification of loaded adjuvant, average particle size, polydispersity and morphology (Figure 3B). Blank and adjuvant-loaded PSs demonstrated sizes between 140 and 155 nm with a polydispersity index (PDI) of less than 0.3. Cryogenic transmission electron microscopy (Cryo-TEM) images confirmed the vesicular morphology of self-assembled PSs (Figure 3C). Small-angle X-ray scattering (SAXS) profiles of Blank-PSs and dual adjuvant-loaded PSs ((CL075+cGAMP)-PS) optimally fits a vesicle model, which further verifies the vesicle morphology of nanocarrier formulations (Figure 3D). The loading of agonists into PSs was generally observed with encapsulation efficiencies of >80% for CL075 or ∼6% for cGAMP (Figure 3E). *in vitro* release of CL075 and cGAMP from PSs was performed over 2 weeks (14 days) in phosphate-buffered saline (pH 7.4) at 37 LJ, with cumulative release ranging from 20 – 60 % (Figure 3F). The highly hydrophilic nature of cGAMP might have contributed to its lower encapsulation efficiency and greater diffusion from PS as compared to hydrophobic CL075 that is trapped inside the PS bilayer membrane.

**Figure 3.**
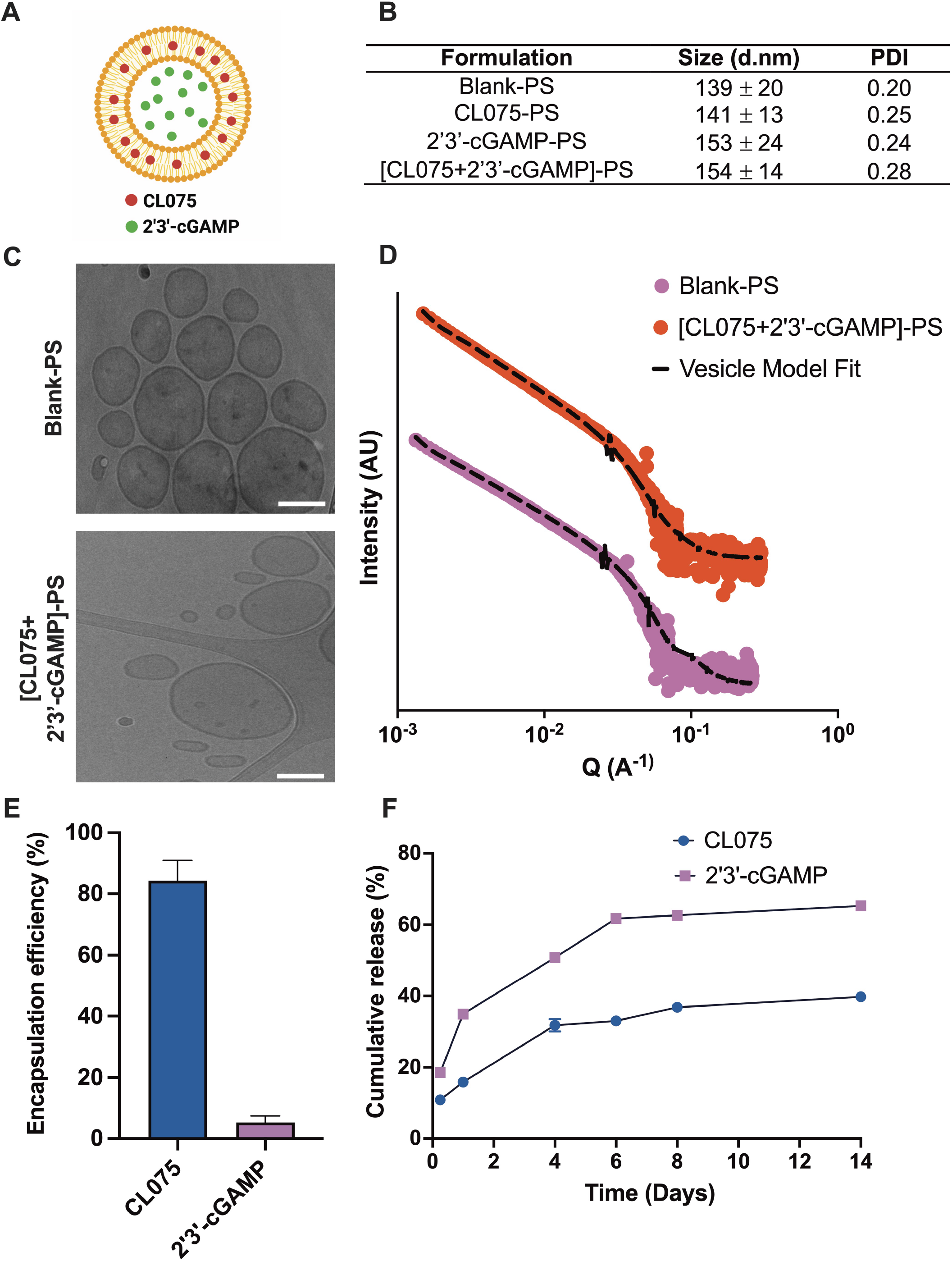
cGAMP and CL075 encapsulation and characterization of adjuvant-loaded PEG-*b*-PPS polymersomes. (A) Schematic showing adjuvant-loaded-PEG-*b*-PPS-PSs, with hydrophobic CL075 in the PS bilayer membrane and hydrophilic cGAMP in the aqueous core. (B) Dynamic light scattering analysis of blank and adjuvant-loaded PS formulations. Size (d.nm) and polydispersity index (PDI) was reported as mean ± SD (n = 3), (C) Representative cryo-TEM images of Blank-PS and dual adjuvant-loaded PS ((CL075+cGAMP)-PS). TEM images were acquired at ×10,000 magnification (scale = 100 nm) (D) Small-angle X-ray scattering (SAXS) scattering profile and model fits for Blank-PS and dual adjuvant-loaded PS ((CL075+cGAMP)-PS). The scattering profile and vesicle-model fit are represented as solid dots and dotted lines, respectively. (E) Encapsulation of CL075 and cGAMP in PS. The encapsulation of adjuvants was measured after purifying PS with size exclusion chromatography. Encapsulation efficiency (%) was reported as mean ± SD (n = 3). (F) *in vitro* release of CL075 and 2’3’-cGAMP from PS over 2 weeks (14 days). Release studies were performed in phosphate-buffered saline (pH 7.4) at 37. Cumulative release (%) was reported as mean ± SD (n = 3).

### Admixture of cGAMP-PS and CL075-PS enhances Th1-polarized Ag-specific adaptive responses in young mice

Previously, we have demonstrated that cGAMP is a promising adjuvant candidate for early life immunization^13^. We have also demonstrated that optimization of vaccine formulation and delivery route can enhance the adjuvanticity potential of TLR7/8 agonists (e.g., CL075)^1, 3^, specifically by encapsulation within PEG-*b*-PPS nanocarriers^12^. Our team has validated PEG-*b*-PPS PSs for nontoxic adjuvant delivery to mice with enhanced targeted uptake by DCs and monocytes^10–12^. To assess whether the synergy observed for the combination of cGAMP and CL075 *in vitro*, was also evident *in vivo*, we next evaluated adjuvant/s loaded PS nanocarriers in an infant mouse vaccination model. Infant mice were immunized intramuscularly (i.m.) by cGAMP (1 µg) and/or CL075 (164 µM) either alone or in an encapsulated or co-encapsulated formulation (Table S1). rHA (Flublok^®^) was used as a model antigen to assess the adaptive immune response. On day of life (DOL) 21, which was 7 days after the boost, serum was collected and analyzed for a rHA specific humoral responses (Figure 4A). Compared to the control groups (PBS, rHA or rHA + Blank-PSs), both cGAMP-PSs and the PS admixture (cGAMP-PSs + CL075-PSs) immunized groups showed significant induction of rHA-specific total IgG (Figure 4B), IgG1 (Figure 4C) and IgG2c (Figure 4D). The co-encapsulated (cGAMP + CL075)-PS formulation did not demonstrate any superior humoral effect with rHA (Figure S1A-C). Next, we focused on the adaptive compartment of vaccine-induced immunity by harvesting splenocytes on DOL26 (12 days after boost) and restimulated the cell suspension with vaccinal antigen (Flublok^®^) for 18 h. The importance of Flublok^®^-specific CD4^+^ T cells, especially the Th1 subset, and CD8^+^ T cells for the development of a proper anti-influenza immunity is now well-established ^27, 28^. Of note, the admixture (cGAMP-PS + CL075-PS) formulation induced a more robust Th1 polarization upon memory recall in the infant setting (Figure 4E-G), characterized by the significant release of IFNγ (Figure 4E), TNF (Figure 4F) and IL-2 (Figure G) compared to control groups as well as cGAMP-PS immunized groups. Co-encapsulated formulation ((cGAMP + CL075)-PS) demonstrated inferior Th1 polarization upon vaccination (Figure 5 & Figure S1D-F). A similar trend was observed in rHA-specific CD8^+^ T cell compartment, where admixture (cGAMP-PS + CL075-PS) triggered IFNγ^+^ release (Figure S2), which has already been proven important in viral clearance upon influenza infection^29–32^. Furthermore, Flublok^®^ stimulation facilitated the triggering of a population of monofunctional IFNγ^+^ TNF^-^ IL-2^-^ CD4^+^ T cells in the co-encapsulated formulation immunized group (Figure S3B).

**Figure 4.**
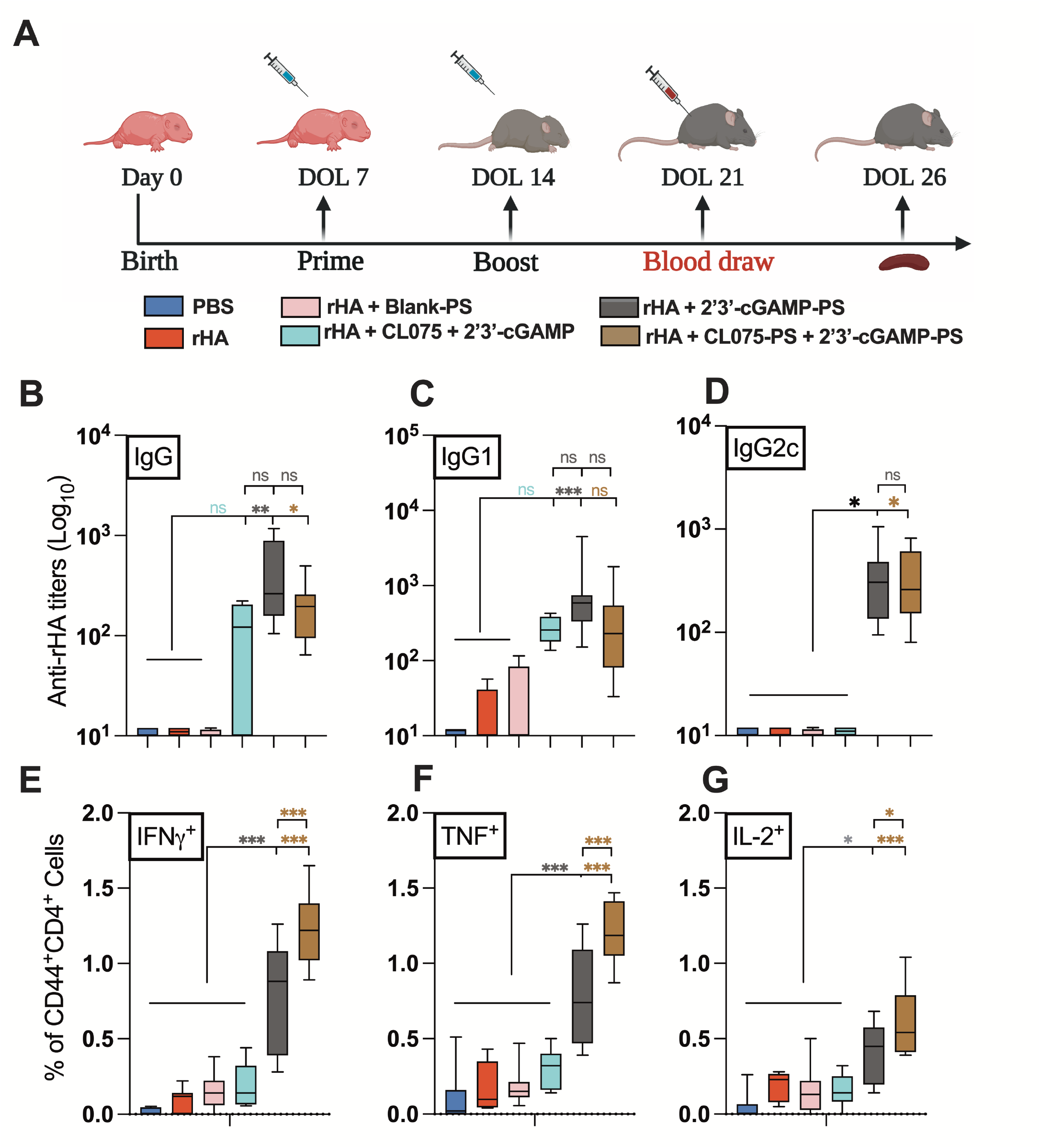
Individually encapsulated cGAMP and admixed cGAMP and CL075 PSs enhanced rHA-specific neonatal humoral and cell mediated immune responses. (A) Infant C57BL/6 mice were immunized i.m. on DOL (day of life) 7 and 14. All groups received 1 μg of each of the Flublok^®^ quadrivalent antigens (rHA), except the PBS group. rHA was given alone or in combination with cGAMP (1 μg) and CL075 (164 μM) delivered in add mixture or single-loaded or dual-loaded PEG-b-PPS nanocarriers. (B) Antibody titers for rHA-specific IgG (B), IgG1(C) and IgG2c (D) were determined by ELISA in serum samples collected at DOL 21. (E-G) Murine CD4^+^ T cell responses after rHA stimulation. Splenocytes from rHA and adjuvanted vaccinates were isolated, stimulated with 10 μg/mL of Flublok^®^ along with CD28 (1 μg/mL) and CD49d (1 μg/mL) for 12 h followed by 6 h of BFA stimulation to block the extracellular cytokine secretion. After stimulation, cells were harvested, stained (intracellular cytokine staining) and analyzed by flow cytometry. Plots were gated on CD44^+^ CD4^+^ lymphocytes and analyzed for all combinations of simultaneous IFNγ, TNF and IL-2 productivity. Statistical comparison employed test one-way ANOVA; \**p* < 0.033, \*\**p* < 0.002, \*\*\**p* < 0.001 (n = 5 – 12 per group), with comparison to PBS, rHA and/or mock loaded PS nanocarriers control groups. Study was inclusive of two independent repeats.

**Figure 5.**
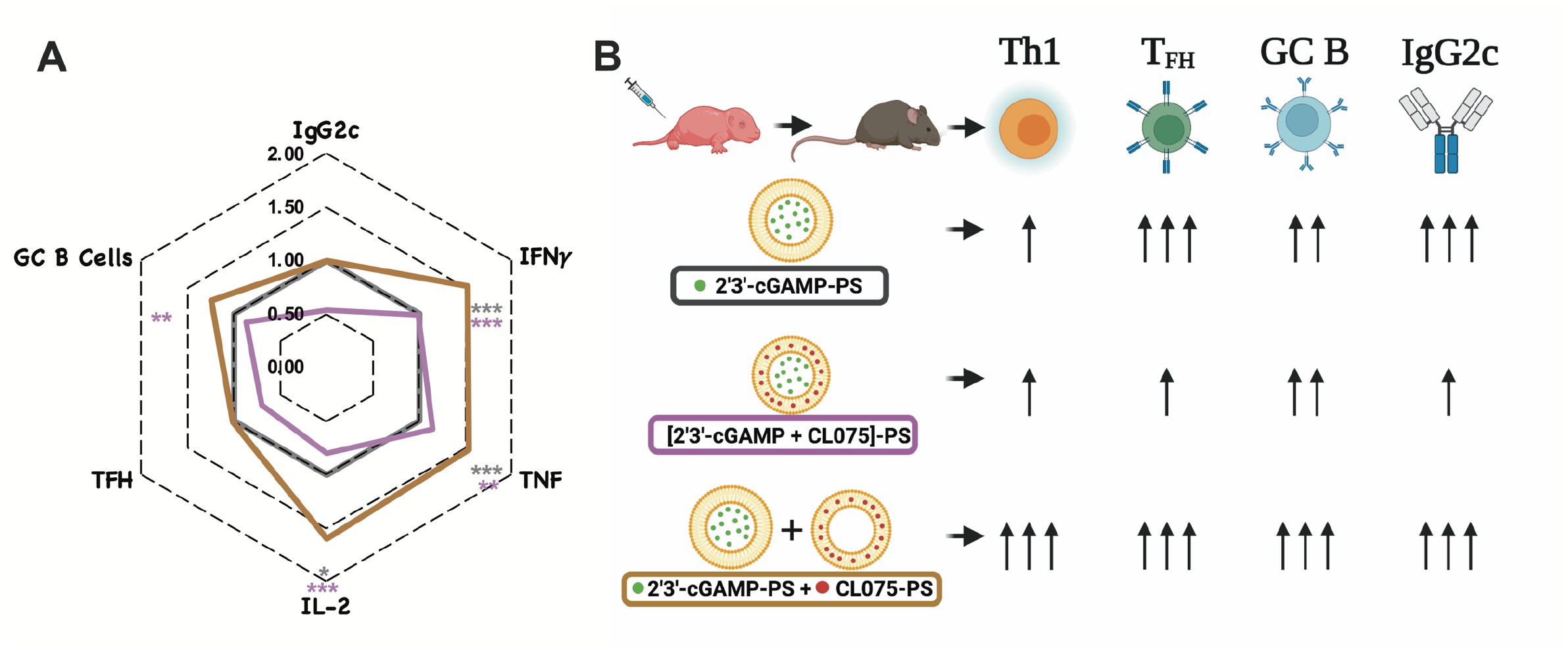
Adjuvanted nanocarrier formulation skews neonatal immunity toward a Type 1 response. Together, cGAMP and CL075 encapsulating PSs alter rHA-specific Th1 polarization and overcome the inability of the infant immune system to mount a type 1 immunity and tune the degree of immune response enhancement. (A) The magnitude of rHA-specific IgG2c titers (at DOL 21), Th1 polarization, T_FH_ and GC B cell responses (DOL26) was shown in a radar plot as a fold-change over cGAMP-PS immunized group (black line). Statistical comparison employed test one-way ANOVA; \**p* < 0.033, \*\**p* < 0.002, \*\*\**p* < 0.001 (n = 5 – 12 per group). Study was inclusive of two independent repeats. (B) Tunable aspects of different formulations when small molecule agonists are encapsulated in PEG-*b*-PPS nanocarriers.

To decipher the immunomodulatory effect of the admixture on germinal centers (GCs), we investigated the accumulation of T_FH_ cells and GC B cells, plasmablasts and plasma cells by flow cytometry in the inguinal and popliteal draining lymph nodes (dLNs) on 12 days after the boost (Figure 5 & Figure S4). Consistent with previous observations that TLR8 signaling evokes T_FH_ cell differentiation *ex vivo*^33, 34^ and that cGAMP with alum^13^ or cGAMP loaded virus-like particles^35^ induce T_FH_ cells in dLNs of adult mice, cGAMP-PSs induced CD4^+^ T_FH_ responses, as reflected by an increased in CD3^+^CD4^+^PD-1^+^CXCR5^+^ T cells (Figure S4C). Despite being the inferior Th1 inducer *in vivo*, the co-encapsulated formulation triggered similar T_FH_ responses when compared with the admixture formulation (Figure S1G). This may explain why cGAMP-PSs and co-encapsulated formulations act as an effective inducer of Flublok^®^-specific humoral responses, since T_FH_ differentiation has proven beneficial for inducing strong humoral immune responses^33, 34^. Furthermore, the admixture (cGAMP-PS + CL075-PS) formulation triggers the frequency of GC B cells in the dLN compartment over the blank-PS control group (Figure S4D). The magnitude of rHA-specific IgG2c titers (at DOL 22), Th1 polarization, T_FH_ and GC B cell responses (DOL26) is depicted in a radar plot as a fold-change over the cGAMP-PS immunized group (black line) (Figure 5A). Fascinatingly, due to lymphadenopathy (unpublished observations), cGAMP-PS immunized group showed a remarkable accumulation of the total number of T_FH_ cells, GC B cells, plasmablasts and plasma cells (Figure S4G-J). Together, cGAMP and CL075 encapsulating PSs alter rHA specific Th1 polarization, overcome the inability of the infant immune system to mount a type 1 immune response, and allow tuning of the degree of immune response enhancement (Figure. 5B).

## CONCLUSION

Newborns and young infants demonstrate distinct immune ontogeny including a reduced ability to drive IFNγ-driven type 1 immunity, which in turn leads to a higher risk of infections with intracellular pathogens and reduced vaccine efficacy ^36, 37^. Although there is no comprehensive consensus on whether and how *in vitro* models can predict the *in vivo* effect of candidate adjuvants, the use of DCs has some advantages for assessing adjuvant activity *in vitro* ^37–40^. For example, *in vitro* and pre-clinical *in vivo* studies have shown that targeting endosomal and cytosolic PRRs, such as TLR7/8 ^41–47^, potently activates human newborn leukocytes and markedly enhances vaccine efficacy in neonatal non-human primates ^43^. In the present work, by combining an *in vitro* study of newborn DC activation and murine *in vivo* immunization model and taking advantage of a delivery systems such as PEG-*b*-PPS PSs, we demonstrated the potential immunization benefit of dual STING and TLR7/8 agonist adjuvant activation in the early life setting. Specifically, using an admixture of cGAMP-PS and CL075-PS as an adjuvantation strategy for early-life immunization, we were able to induce cardinal features of type 1 immunity: 1) IFNγ production by antigen-specific CD4^+^ T cells and 2) relatively high titers of antigen-specific IgG2c. As IFNγ promotes isotype switching toward IgG2a/c *in vivo* ^48^, these two effects are likely linked.

Interestingly, in our study, we did not observe any benefit from co-encapsulation of cGAMP and CL075 within PS as compared with individual encapsulation and admixed agonist PSs. This may be due to the faster release kinetics of cGAMP from encapsulated PS compared to CL075, due to its hydrophilic nature. Alternatively, our admixed approach may simply generate distinct heterogenous activated APC populations. In the case of co-encapsulation only one activated APC population would be anticipated – i.e., those receiving both cGAMP and CL075 while the admixture may generate 3 distinct populations: 1) APCs receiving cGAMP, 2) APCs receiving CL075 and 2) APCs receiving both cGAMP and CL075. The latter may manifest distinct immunostimulation of cGAMP and CL075.

Our study also expands upon prior materials science work ^49^, where the STING agonist 3’3’-cGAMP and/or the TLR8/7 agonist R848 were encapsulated using acetylated dextran (Ace-DEX), and co-encapsulation of 3’3’-cGAMP and R848 trended towards enhanced ovalbumin-specific humoral immunity (IgG and IgG1) as compared to the individually encapsulated adjuvants. In contrast to our studies, the release kinetics of 3’3’-cGAMP or R848 were similar to co-encapsulated Ace-DEX ^49^. Furthermore, our use of multi-parameter flow cytometry on both splenocytes and dLN derived cells demonstrated a unique age-specific response *in vivo* to licensed influenza vaccines. Such responses cannot be fully replicated via the use of an ovalbumin-specific response as quantified by ELISpot assay or by ELISA using supernatant from protein-stimulated murine splenocytes ^49^.

Overall, our study features several strengths, including 1) use of primary murine and human DCs to model individual and combined adjuvant effects *in vitro*, 2) assessment and demonstration of age-specific synergy of combined engagement of STING and TLR7/8 to drive robust activation of neonatal DCs, 3) mechanistic demonstration of a precision expansion of early life Th1 cells via the IL-12/IFNY axis, and 4) assessment of the individual and combined effects of the adjuvants in mice *in vivo*. Our study also has several limitations, including 1) the potential effects of cGAMP-PS and CL075-PS on GCs are intriguing, but we might have captured delayed GC B cells and T_FH_ responses^50^, 2) although our studies demonstrated robust enhancement of immunogenicity with cGAMP-PSs and CL075-PSs together, future functional studies (e.g., pathogen challenge) are required to assess the efficacy of the this adjuvantation approach, and 3) due to species specificity, *in vivo* results in mice may not accurately reflect *in vivo* effects in humans, and would be further supported by porcine and or non-human primate studies.

The flexibility and adaptability of the PS technology can be leveraged to fine-tune vaccine immunogenicity via targeted antigen or small molecule adjuvant delivery and may represent a multifunctional platform for precision vaccine design in the 21^st^ century. Such an approach must account for tailoring vaccines to vulnerable populations with distinct immunity. For example, receptor-binding domain (RBD) protein and MPLA can be formulated into stable, biologically active PSs, and can induce robust RBD specific humoral and polyfunctional Th1 responses in adult mice ^51^. However, adjuvantation systems such as AS01 and AS02, both consisting of MPLA and the purified plant bark extract/saponin QS21, are components of the *Mosquirix* (RTS,S) malaria vaccine ^52^, which has demonstrated reduced efficacy in the pediatric setting ^52, 53^.

In conclusion, we demonstrate that individually encapsulated and admixed cGAMP-PSs and CL075 PSs shape the quantity and quality of neonatal immune responses, Th1 polarize neonatal rHA-specific humoral and cell-mediated immune responses, presenting a promising adjuvant approach for early life immunization. Since we employed the recombinant hemagglutinin influenza vaccine throughout our work, our results may be applicable to early life influenza immunization. The use of age-specific *in vitro* and *in vivo* modeling may also represent a general strategy for optimizing type 1 immunity towards protein antigens for early life immunization against intracellular pathogens including additional respiratory viruses such as coronaviruses.

## SUPPORTING INFORMATION

**Figure S1. Comparison of rHA specific humoral and cell-mediated responses in between co-encapsulation and individual encapsulation of cGAMP and CL075 in PEG-*b*-PPS nanocarriers.** After immunization of infant C57BL/6 mice i.m. on DOL (day of life) 7 and 14, antibody titers for rHA-specific IgG (A), IgG1(B) and IgG2c (C) were determined by ELISA in serum samples collected at DOL 21. (D-F) Splenic CD4^+^ T cell responses after rHA stimulation. (G-I) Total number of T_FH_ cells (CD3^+^CD4^+^PD-1^+^CXCR5^+^), GC B cells (CD3^-^CD19^+^CD95^+^GL7^+^), plasmablasts (CD3^-^ CD19^+^ CD138^+^) in DLN. Statistical comparison employed test one-way ANOVA; \**p* < 0.033, \*\**p* < 0.002, \*\*\**p* < 0.001 (n = 5 – 7 per group). Study was inclusive of two independent repeats.

**Figure S2. Gating strategy to identify rHA-specific CD8^+^ T cells.** (A) Splenocytes were isolated from immunized mice following 12 days booster. Antigen-specific T cell responses following rHA stimulation were defined as CD3^+^CD8^+^CD44^high^ IFNγ^+^ using FlowJo software, v.10.8.1. Shown was an example of the hierarchical gating strategy leading to the identification of live, singlet, CD3^+^, CD4^+^, CD44^high^ and IFNγ^+^ T cells in different immunize groups. (B) Splenic CD8^+^ IFNγ^+^ T cell signature was analyzed from indicated immunized groups after 12 days booster. Statistical comparison employed test one-way ANOVA; *p < 0.033, **p < 0.002, ***p < 0.001 (n = 5 – 12 per group). Study was inclusive of two independent repeats.

**Figure S3. Gating strategy to identify rHA-specific CD4^+^ T cells and it’s multifunctionality.** (A) Splenocytes were isolated from immunized mice following 12 days booster. Antigen-specific T cell responses following rHA stimulation were defined as CD3^+^CD4^+^CD44^high^ cytokine^+^ using FlowJo software, v.10.8.1. Shown was an example of the hierarchical gating strategy leading to the identification of live, singlet, CD3^+^, CD4^+^, CD44^high^ and cytokine^+^ T cells with an example for IFNγ, TNF and IL-2 response in different immunize groups. (B) Multifunctionality of the CD4^+^ IFNγ^+^ T cells cytokine response was analyzed from indicated immunized groups after 12 days booster. Pie chart represent the fraction of the total CD4^+^ IFNγ^+^ cytokines response comprising any combination of TNF and IL-2 production after Flublok^®^ stimulation in different immunized groups. Beneath pie chart, bar graph represents the frequencies of multifunctional T cells in CD4^+^ IFNγ^+^ T cell compartment. Statistical comparison employed test one-way ANOVA; *p < 0.033, **p < 0.002, ***p < 0.001 (n = 5 – 12 per group). Study was inclusive of two independent repeats.

**Figure S4. cGAMP and CL075 encapsulating PS promotes T_FH_ and B cell responses in draining lymph nodes (DLN).** (A) Shown was an example of the hierarchical gating strategy leading to the identification of T_FH_ cells (CD3^+^CD4^+^PD-1^+^CXCR5^+^), GC B cells (CD3^-^CD19^+^CD95^+^GL7^+^), plasmablasts (CD3^-^CD19^+^ CD138^+^) and plasma cells (CD3^-^CD19^-^CD138^+^) are shown. (B) Infant C57BL/6 were immunized i.m. as described in Figure 4. Lymphocytes from DLN (both popliteal and inguinal) of immunized mice were harvested at DOL 26 for FACS analysis. (C) % of T_FH_ cells, (D) % of GC B cells, (E) % of Plasma cells and (F) % of plasmablasts among live cells. (G-J) Total number of immune subsets in DLN. Statistical comparison employed test one-way ANOVA; *p < 0.033, **p < 0.002, ***p < 0.001 (n = 5 – 7 per group), with comparison to rHA and mock loaded PEG-b-PPS nanocarriers control groups. Study was inclusive of two independent repeats.

**Figure S5. Comparison of rHA specific humoral and cell mediated responses in between different formulations.** After immunization of infant C57BL/6 mice i.m. on DOL (day of life) 7 and 14, antibody titers for rHA-specific IgG (A), IgG1(B) and IgG2c (C) were determined by ELISA in serum samples collected at DOL 21. (D-F) Splenic CD4^+^ T cell responses after rHA stimulation was determined by flow cytometry. Statistical comparison employed test one-way ANOVA; *p < 0.033, **p < 0.002, ***p < 0.001 (n = 5 – 12 per group). Study was inclusive of two independent repeats.

**Table S1. Vaccine formulations for *in vivo* study.**

**Table S2. Panels and regents for flow cytometry assay.**

## MATERIALS AND METHODS

### Animals and ethical declaration

All experiments were conducted in accordance with relevant institutional and national guidelines, regulations, and approvals. All experiments involving animals were approved by the Institutional Animal Care and Use Committees (IACUC) of Boston Children’s Hospital and Harvard Medical School (protocol numbers 19-02-3792R and 19-02-3897R). C57BL/6 mice (either 6-8 weeks or pregnant) were obtained from Jackson Laboratories (Bar Harbor, ME) and housed in specific pathogen-free conditions in the animal research facilities at Boston Children’s Hospital. Cages checked daily to assess the presence of pups. Discovery of a new litter was recorded as DOL 0. Both male and female pups were used for neonatal experiments in a littermate-controlled specific manner. CO_2_ was used as the primary euthanasia method, with cervical dislocation as a secondary physical method to ensure death.

Non-identifiable human cord blood samples were collected with approval from the Ethics Committee of The Brigham & Women’s Hospital, Boston, MA (Institutional Review Board (IRB) protocol number 2000-P-000117) and Beth Israel Deaconess Medical Center Boston, MA (IRB protocol number 2011P-000118). Blood samples from adult volunteers were collected after written informed consent with approval from the Ethics Committee of Boston Children’s Hospital, Boston, MA (IRB protocol number X07-05-0223).

### Murine Bone Marrow-Derived Dendritic Cells (BMDCs) assay

BMDCs were generated from newborn (5 – 7 days old) and adult (6 – 12 weeks old) C57BL/6 mice with an adaptation of previously described methods ^13, 54, 55^. Briefly, mice were sacrificed followed by the surgical removal of femurs and tibiae. Bones were surgically cleaned from surrounding tissue, extremities of tibiae and femurs were trimmed with sterile scissors and bone marrow was flushed through a 70 μm nylon mesh strainer (Corning Life Sciences). Cell number and viability were determined by the trypan blue exclusion method. Whole bone marrow cells were plated into non-tissue culture-treated 100 mm Petri dishes (Corning Life Sciences) at a density of 0.3 x 10^6^ cells/mL in 10 mL total volume/plate of complete culture medium (RPMI 1640 plus 10% heat-inactivated fetal bovine serum (FBS, GE Healthcare HyClone), 50 μM 2-mercaptoethanol, 2 mM L-glutamine, 100 U/mL penicillin/streptomycin (Gibco ThermoFisher Scientific) supplemented with 20 ng/ml of recombinant murine GM-CSF (rmGM-CSF, R&D systems). Plates were incubated in humidified atmosphere at 37°C, 5% CO_2_ for 6 days, with one supplement of 10 mL of complete culture medium and rmGM-CSF on day 3. On day 6, non-adherent and loosely adherent cells were harvested by washing the plate gently with culture medium. Adherent cells were discarded.

For stimulation experiments, immature BMDCs generated from 7 day old and adult mice were plated in round bottom 96-wells non tissue culture-treated plates at a density of 10^5^ cells/well in 200 μL of fresh complete culture medium with rmGM-CSF as described above, with the appropriate stimuli. Cells were incubated at 37°C for 20-24 h, then supernatant harvested and TNF and IL-12p70 concentrations were measured by ELISA (R&D Systems). IFNβ was measured with a bioluminescent ELISA kit (LumiKine, Invivogen). For experiments involving blocking antibodies (Abs), BMDCs were pre-incubated for 20 minutes at 37 °C with anti-mouse IFNAR1 (clone MAR1-5A3, 10 μg/mL, Biolegend), anti-mouse IL-12p40 or anti-mouse IL-12p70 Abs or an isotype control before stimulation.

### Human Monocyte Derived Dendritic Cells (MoDCs) assay

Following local IRB-approved protocols, peripheral blood was collected from healthy adult volunteers, while human newborn cord blood was collected immediately after Cesarean section delivery of the placenta. Births to known HIV-positive mothers were excluded. Human blood was anti-coagulated with 20 units/mL pyrogen-free sodium heparin (American Pharmaceutical Partners, Inc.; Schaumberg, IL). All blood products were kept at room temperature and processed within 4 h from collection. Primary human PBMCs and CBMCs were isolated from fresh blood via Ficoll gradient separation.

Monocytes were isolated from PBMC and CBMC fractions via positive selection by magnetic microbeads according to the manufacturer’s instructions (Miltenyi Biotec, Auburn, CA) using CD14 as a pan marker. Isolated monocytes were cultured in tissue culture dishes at 0.4 × 10^6^ cells/mL in RPMI 1640 media containing fresh 10% autologous plasma, supplemented with recombinant human IL-4 (50 ng/mL) and recombinant human GM-CSF (100 ng/mL) (R&D Systems, Minneapolis, MN) with one additional supplement of fresh media and cytokines at day 3 of culture as previously described ^12, 44^. After 6 days, immature MoDCs were harvested by gently pipetting the loosely adherent fraction, before being re-plated (10^5^ cells/ well) in 96-well flat-bottom plates in the presence or absence of TLRs, and/or 2% Glycerol, and/or sterile PBS. Plates were then incubated for 18-24 h at 37°C in a humidified incubator at 5% CO_2_. After this stimulation supernatants were harvested and processed for further functional assays.

### Human newborn cord blood mononuclear cells (CBMCs) stimulation

Human newborn cord blood mononuclear cells (CBMCs), containing a T cell compartment largely composed of naïve T cells, were plated at 0.1 x 10^6^ cells/well in 96-well tissue culture plates and stimulated with aqueous formulations of indicated compounds, in the presence of the polyclonal T cell activator αCD3 (Plate-bound, 5 μg/mL) for 96 h. Culture supernatants were harvested and analyzed for IFNγ induction using via IFNγ ELISA (InvivoGen). For some experiments, CBMCs were stimulated with agonists alone or in combination with cytokine blocker for the last 6 h. After stimulation, cells were harvested, stained with surface markers followed by intracellular cytokine markers as described in Table S2 as previously described ^13^. Stained cells were analyzed by flow cytometer to quantify the percentage of T cells which producing IL-4, IL-17 and IFNγ.

### Nanocarriers formulations

A range of PS formulations were prepared using PEG-*b*-PPS block copolymer, including blank (unloaded) PS and PS loaded with CL075, cGAMP and CL075+cGAMP (Table S1). PS nanocarriers were prepared by flash nanoprecipitation technique utilizing a confined-impingement (CIJ) mixer as published earlier ^7^. Here, the organic phase was prepared by dissolving 20 mg PEG-*b*-PPS polymer in 500 μL of tetrahydrofuran (THF) and 200 μg of CL075 while the aqueous phase was prepared by dissolving 2 mg of cGAMP in phosphate-buffered saline (PBS, pH 7.4). Organic and aqueous phases were separately loaded into syringes and impinged against each other into 2 mL aqueous reservoir containing PBS. THF was removed from the formulation using overnight vacuum desiccation. PS were then purified to remove unencapsulated CL075 and cGAMP using Sephadex LH-20 and Sepharose CL-6B columns, respectively. For Blank PS, organic and aqueous phases were added without adjuvants whereas single adjuvant-loaded PS contained the adjuvant of interest either in organic or aqueous phase.

### Size and morphological characterization of polymersomes

Dynamic light scattering (DLS) analysis was performed using a Zetasizer Nano-ZS (Malvern Instruments, UK) to measure the size of PS. All samples were diluted 1 in 1000 with PBS prior to the analysis.

For cryogenic transmission electron microscopy (Cryo-TEM) studies, Pelco easiGlow glow discharger (Ted Pella) was used to glow discharge the lacey carbon Cu grids (200 mesh) at 15 mA atmosphere plasma. The grids were applied with 4 μL volume of PS sample, blotted for 5 s and plunged into liquid ethane within a FEI Vitrobot Mark III plunge freezing instrument (Thermo Fisher Scientific). Grids were stored in liquid nitrogen. Grids were viewed on a JOEL JEM1230 LaB6 emission TEM (JOEL USA) at 100 keV and micrographs were obtained using a Gatan Orius SC1000 CCD camera Model 831 (Gatan).

Small-angle x-ray scattering (SAXS) studies were performed at the DuPont-Northwestern-Dow Collaborative Access Team (DND-CAT) beamline at Argonne National Laboratory’s Advanced Photon Source (Argonne, IL, USA). Collimated X-rays with 10 keV (wavelength λ = 1.24 Å) was utilized to measure samples. Sample measurement was performed in the q-range 0.001 to 0.5 Å^-1^, which was calibrated using silver behenate. The final scattering data of samples was obtained using PRIMUS 2.8.2 software after subtracting solvent buffer scattering from sample scattering. The morphologic characteristics of PS samples were confirmed after fitting vesicle model with sample scattering data in SasView 4.0 software.

Encapsulation of CL075 and cGAMP in PS was measured after purifying PS with size exclusion chromatography and efficiency reported as mean ± SD (n = 3). The purified adjuvant-loaded PS samples obtained as mentioned above were lyophilized to rupture the nanocarrier structure. For CL075 samples, the lyophilized samples were dissolved in HPLC grade DMF and encapsulation was measured by means of HPLC-UV/fluorescence against known standards with a dimethylformamide mobile phase, as previously described ^12^. Each formulation batch was stored at -20°C until day of use. For cGAMP, lyophilized samples were added with methanol to precipitate PEG-*b*-PPS block copolymer and dissolve cGAMP. The polymer precipitate was removed via centrifugation and supernatants containing cGAMP were quantified using LC-MS method as previously described^56^.

### Immunization and rHA specific antibody quantification

For neonatal mouse studies, 6-day old pups were toe clipped for individual recognition, 7-day old C57BL/6 mice were immunized with a prime-boost schedule (two injections, each one week apart, for newborn mice at DOL 7 and 14). Neonatal mice were immunized i.m. in the posterior thigh with 50 μL of total vaccine dose, separated across both hind legs (25 μL per leg) according to Table S1. Each 50 µL of vaccine dose was included 1 µg of each of the following recombinant influenza virus hemagglutinins (rHA): A/Hawaii/70/2019 (H1N1), A/Minnesota/41/2019 (an A/Hong Kong/45/2019-like virus) (H3N2), B/Washington/02/2019 and B/Phuket/3073/2013 contained in the 2020-2021 formulation of the Flublok^®^ vaccine (Protein Sciences Corp.). Serum was harvested 7 days following boost (day of life 21) via retroorbital bleed and anti-rHA serum total IgG titers, IgG1 and IgG2c were measured by ELISA. During the immunization or retroorbital bleed, mice were anaesthetized with isoflurane.

For anti-rHA ELISAs, CoStar 96 well high-binding plates (Corning, Corning, NY) were coated with 1 µg/mL rHA in carbonate buffer pH 9.6, incubated overnight at 4°C, washed 3x with wash buffer (KPL 10X Phosphate Buffered Saline with 0.05% Tween 20 (Fisher Scientific)) and blocked with Superblock (ScyTek) for 1 h at room temperature (RT). Then, sera from vaccinated mice were added with an initial dilution of 1:100 and 1:2 serial dilutions in EIA buffer (PBS + BSA 1% + Tween 20 0.1% + Heat inactivated FBS 10%) and incubated for 2h at RT. Plates were then washed 3x and incubated for 1h at RT with HRP-conjugated anti-mouse IgG, IgG1 or IgG2c (Southern Biotech). At the end of the incubation plates were washed 5x and developed with BD OptEIA TMB substrate reagent set (BD, San Jose, CA) for 10 minutes, then stopped with 1 M H_2_SO_4_. The optical density was measured at 450 nm with SpectraMax ID3 microplate reader with SoftMax Pro Version 5 (both from Molecular Devices) and either titers or concentrations were calculated using as cutoff three times the optical density of the background.

### Cell mediated immune responses *ex vivo*

12 days post booster-immunization (DOL 26), murine spleens were collected in RPMI 1640 media containing 10% heat-inactivated FBS. For analysis of single-cell suspensions, spleens were mechanically and aseptically dissociated with the back of a syringe plunger and filtered through a 70-μm cell strainer and the dissociated tissue was collected in RPMI 1640 media. After centrifugation (500 x g, 10 mins, RT), cells were treated with 1 mL ACK lysis buffer (Gibco, Waltham, MA) for 2 min at RT to lyse red blood cells. Cells were washed immediately with RPMI 1640, passed through 70-μm cell strainer, and suspended in RPMI 1640 media (supplemented with 10% heat-inactivated FBS). Splenocytes were plated at a density of up to 2 × 10^6^ cells/well in a 96-well U-bottom plate and stimulated with 10 μg/mL of rHA (Flublok^®^, 2020-21 formulation) in T cell media. T cell media consists of RPMI 1640 (Gibco, Waltham, MA) supplemented with 10% heat-inactivated FBS (Cytiva HyClone, Fischer Scientific), 100 U/mL Penicillin and 100 mg/mL Streptomycin (Gibco, Waltham, MA), 55 mM 2-mercaptoethanol (Gibco, Waltham, MA), 60 mM Non-essential Amino Acids (Gibco, Waltham, MA), 11 mM HEPES (Gibco, Waltham, MA), and 800 mM L-Glutamine (Gibco, Waltham, MA). In addition to antigen, the stimulation cocktail consisted of 1 μg/mL anti-mouse CD28/49d (BD Biosciences) as a co-stimulant.

For intracellular cytokine staining (ICS), 18 h stimulation was completed in a humidified incubator at 37°C, 5% CO_2_ and 5 μg/mL Brefeldin A (BFA; BioLegend) was added during the last 6 h of stimulation, to block the cytokines production and facilitate optimal intracellular flow cytometry analysis. After 18h of stimulation, cells were washed twice with PBS and blocked with Mouse Fc Block (BD Biosciences) according to the manufacturer’s instructions. After blocking, cells were washed once with PBS and stained with Aqua Live/Dead stain (Life Technologies, Carlsbad, CA) for 15 min at RT. Following two additional PBS washes, cells were resuspended in 100 µL of FACS buffer (PBS supplemented with 0.2% BSA (Sigma-Aldrich)) containing mouse specific cell surface markers for flow cytometry. Markers included anti-mouse CD44 PerCP-Cy5.5, CD3 BV785, CD4 APC/Fire750 and CD8 BUV395. Details of the clone and manufacturer of each marker used in a customized seven colors flow-cytometry panel are documented in Table S2. Cells were incubated with the surface markers for 30 min at 4 LJC. Cells were washed with PBS and fixed/permeabilized by using the BD Cytofix/Cytoperm fixation/permeabilization solution kit according to the manufacturer’s instructions. Cells were washed in 1X perm/wash solution and subjected to intracellular staining (30 min at 4 °C) using a cocktail of the following Abs: anti-mouse IFNγ Alexa Fluor 488, TNF PE Cy7 and IL-2 PE in 1X perm/wash solution. Finally, cells were washed in PBS and fixed in PBS containing 1% paraformaldehyde (Electron Microscopy Sciences, Hatfield, PA) for 20 min at 4 LJC. After two final washes in PBS, the cells were resuspended in PBS and stored at 4 LJC until acquisition. Samples were acquired on a BD LSR II (BD Biosciences; San Jose, CA) configured with blue (488 nm), yellow/green (568 nm), red (640 nm), violet (407 nm), and ultraviolet (355 nm) lasers using standardized good clinical laboratory practice procedures to minimize variability of data generated. Analysis was performed using FlowJo software, v.10.8.1 according to the gating strategy outlined in Figure S3A.

### T_FH_ and B cell responses in Draining lymph nodes

Draining (inguinal and popliteal) lymph nodes (dLNs) from mice were harvested 12 days post-boost (DOL 26) as previously reported^13^. To prepare a single-cell suspension, dLNs were mechanically and aseptically dissociated with the back of a syringe plunger and filtered through a 70-μm cell strainer and collected in RPMI 1640 media. Then, cells were washed and stained with the Abs (described in Supplementary table 2) as previously reported ^57^. The gating strategy was shown in Figure S4A. T_FH_ cells in DLNs were phenotyped as CD3^+^CD4^+^PD-1^+^CXCR5^+^, GC B cells as CD3^-^ CD19^+^CD95^+^GL7^+^, plasmablasts as CD3^-^CD19^+^ CD138^+^ and plasma cells as CD3^-^CD19^-^CD138^+^ by using a customized flow-cytometry panel. Details of the clone, manufacturer and titer of each marker were documented in Table S2.

### Statistical analyses and graphics

Statistical significance and graphic output were generated using GraphPad Prism version 9.3.1 for macOS (GraphPad Software, La Jolla, CA, USA). Data were tested for normality by using the Shapiro-Wilk test. Group comparisons were performed by One-way ANOVA with Dunnett’s multiple comparison post-test or Two-way ANOVA comparing column and row effects. Measurements that failed normality tests were analyzed with Wilcoxon rank-sum tests or a Kruskal-Wallis rank-sum test comparisons within treatments groups. Results were considered significant at *p* values indicated in each figure legend. Synergy was calculated using an adaptation of the Loewe method of additivity as previously described^58^. D values greater than one were considered antagonistic, D values equal to one were considered additive and D values less than one were considered synergistic. Analysis and presentation of flow-cytometric data (Figure S3B) was performed using SPICE (Ver. 6; https://niaid.github.io/spice/) ^59, 60^. Graphics in abstract, Figure 3A, Figure 4A, Figure 5B and Figure S4B were created with BioRender.com.

## Author Contributions

SB and DJD designed the experiments, prepared the figures and wrote the manuscript. FB and CP performed *in vitro* experiments, flow cytometry (human samples) and assisted with data analysis. SB (Bobbala), SY and ES synthesized and characterized adjuvant-loaded nanocarriers. BB precipitated in the animal experiments and organ harvesting. CS, MM and DS helped BB and SB during the preparation of single cell suspension from the harvested organs. SB performed the flow cytometry (mice samples) assay and analyzed the data. MDL, YS, EN and DS were participated in the experiments related with vaccine induced humoral responses. SvH and OL edited and critically reviewed the manuscript. ES, OL, and DJD conceived the project, secured funding and supervised the study.

## Notes

EN, FB, BB, DS, SvH, OL and DJD are named inventors on vaccine adjuvant patents assigned to Boston Children’s Hospital. FB has signed consulting agreements with Merck Sharp & Dohme Corp. (a subsidiary of Merck & Co., Inc.), Sana Biotechnology, Inc., and F. Hoffmann-La Roche Ltd. IZ reports compensation for consulting services with Implicit Biosciences. Rest of the authors do not have any competing interests.

## Funding

DJD’s laboratory is supported by NIH grant 1R21AI137932-01A1. Both DJD and OL’s laboratory are supported by Adjuvant Discovery Program (75N93019C00044), Adjuvant Development Program (272201800047C) and Development of Vaccines for the Treatment of Opioid Use Disorder (272201800047C-P00003-9999). The *Precision Vaccines Program* is supported in part by the Boston Children’s Hospital Department of Pediatrics. CP was supported by the scholarship “J. Miglierina,” Fondazione Comunitaria del Varesotto, Varese, Italy.

## Acknowledgements

We thank the members of the BCH *Precision Vaccines Program* for helpful discussions as well as Drs. Kevin Churchwell, Gary Fleisher, David Williams, and Mr. August Cervini for their support of the *Precision Vaccines Program*. DJD would like to thank Siobhan McHugh, Geneva Boyer, Lucy Conetta and the staff of Lucy’s Daycare, the staff the YMCA of Greater Boston, Bridging Independent Living Together (BILT), Inc., and the Boston Public Schools for childcare and educational support during the COVID-19 pandemic.

## Notes

### Competing Interest Statement

EN, FB, BB, DS, SvH, OL and DJD are named inventors on vaccine adjuvant patents assigned to BCH. FB has signed consulting agreements with Merck Sharp & Dohme Corp. (a subsidiary of Merck & Co., Inc.), Sana Biotechnology, Inc., and F. Hoffmann-La Roche Ltd. IZ reports compensation for consulting services with Implicit Biosciences. Rest of the authors do not have any competing interests.

