## SUPPORTING INFORMATION for "Shaping neonatal immunization by tuning delivery of synergistic adjuvants via nanocarriers"

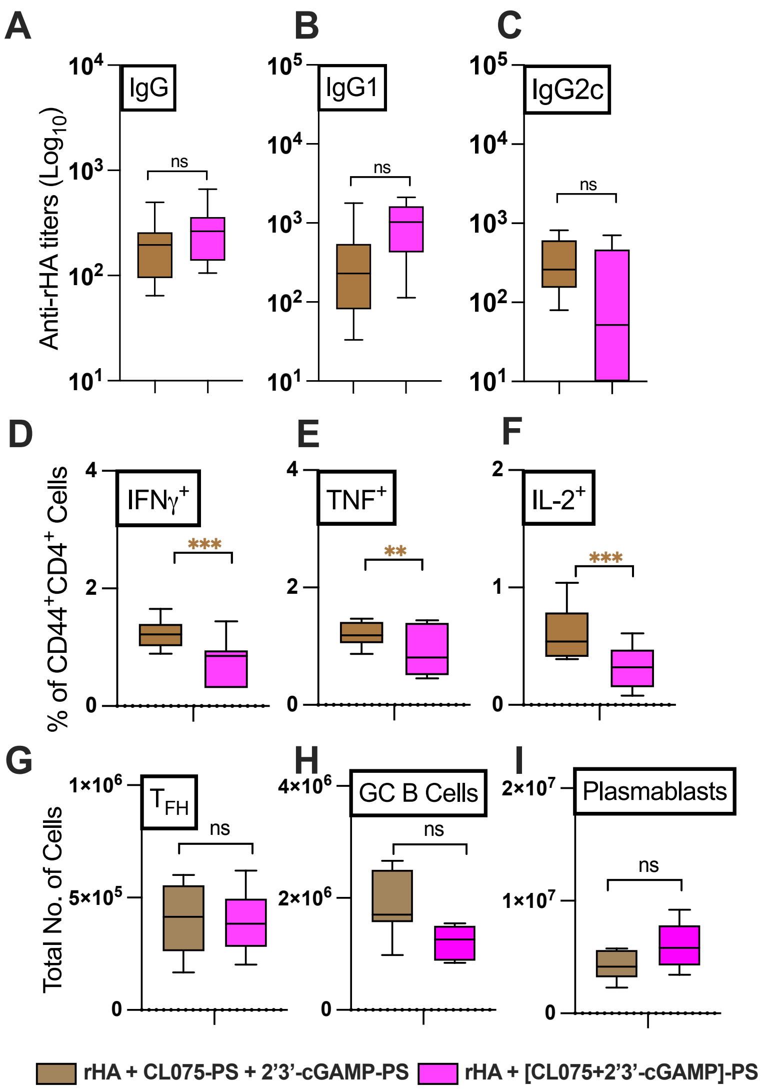


**Figure S1. Comparison of rHA specific humoral and cell-mediated responses in between co-encapsulation and individual encapsulation of cGAMP and CL075 in PEG-*b*-PPS nanocarriers.** After immunization of infant C57BL/6 mice i.m. on DOL (day of life) 7 and 14, antibody titers for rHA-specific IgG (A), IgG1(B) and IgG2c (C) were determined by ELISA in serum samples collected at DOL 21. (D-F) Splenic $CD4^+$ T cell responses after rHA stimulation. (G-I) Total number of $T_{FH}$ cells ($CD3^+CD4^+PD\text{-}1^+CXCR5^+$), GC B cells ($CD3^-CD19^+CD95^+GL7^+$), plasmablasts ($CD3^-CD19^+$ $CD138^+$) in DLN. Statistical comparison employed test one-way ANOVA; **p* < 0.033, ***p* < 0.002, ****p* < 0.001 (n = 5 - 7 per group). Study was inclusive of two independent repeats.

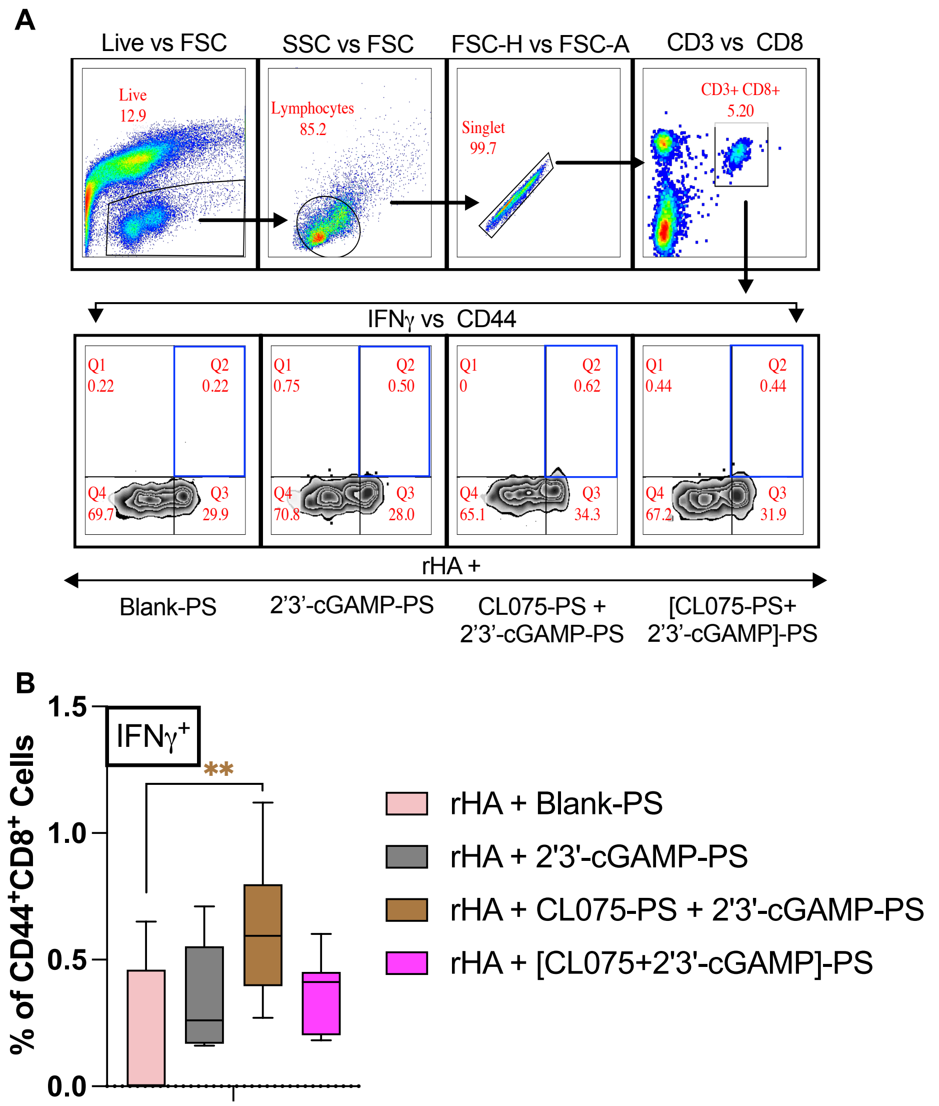


**Figure S2. Gating strategy to identify rHA-specific CD8$^+$ T cells.** (A) Splenocytes were isolated from immunized mice following 12 days booster. Antigen-specific T cell responses following rHA stimulation were defined as CD3$^+$CD8$^+$CD44$^{high}$ IFNg$^+$ using FlowJo software, v.10.8.1. Shown was an example of the hierarchical gating strategy leading to the identification of live, singlet, CD3$^+$, CD4$^+$, CD44$^{high}$ and IFNg$^+$ T cells in different immunize groups. (B) Splenic CD8$^+$ IFNγ$^+$ T cell signature was analyzed from indicated immunized groups after 12 days booster. Statistical comparison employed test one-way ANOVA; *p < 0.033, **p < 0.002, ***p < 0.001 (n = 5 - 12 per group). Study was inclusive of two independent repeats.

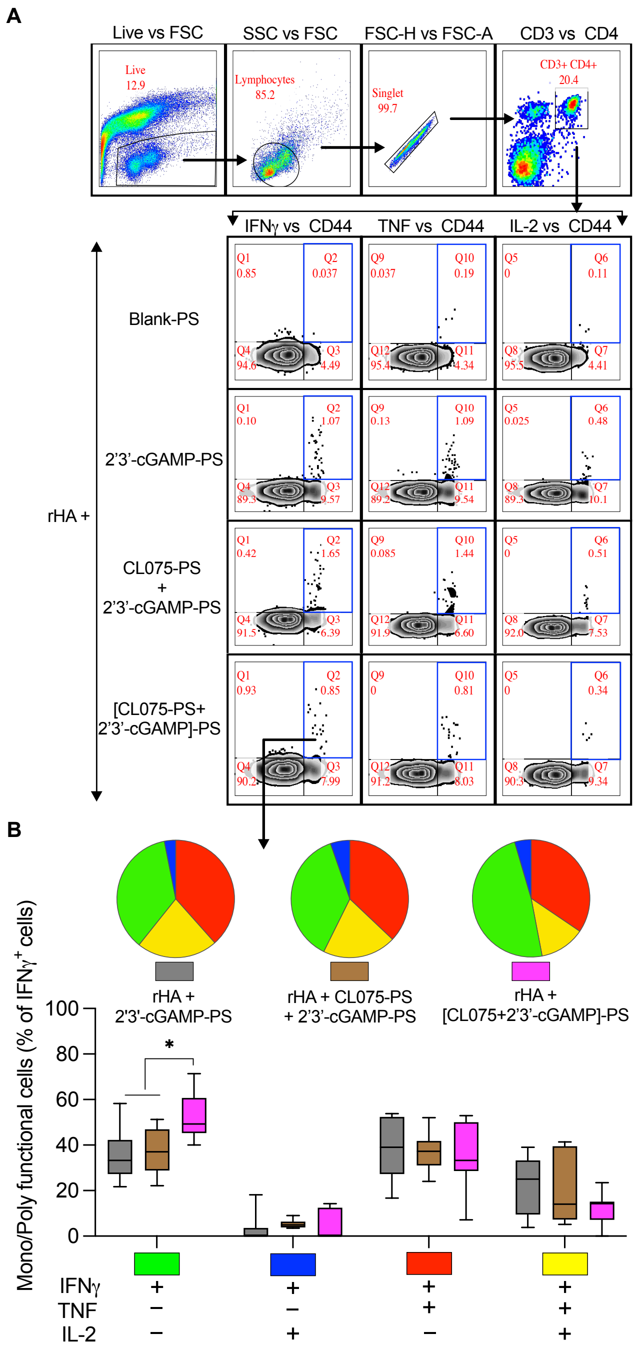


**Figure S3. Gating strategy to identify rHA-specific CD4$^+$ T cells and it's multifunctionality.** (A) Splenocytes were isolated from immunized mice following 12 days booster. Antigen-specific T cell responses following rHA stimulation were defined as CD3$^+$CD4$^+$CD44$^{high}$ cytokine$^+$ using FlowJo software, v.10.8.1. Shown was an example of the hierarchical gating strategy leading to the identification of live, singlet, CD3$^+$, CD4$^+$, CD44$^{high}$ and cytokine$^+$ T cells with an example for IFNγ, TNF and IL-2 response in different immunize groups. (B) Multifunctionality of the CD4$^+$ IFNγ$^+$ T cells cytokine response was analyzed from indicated immunized groups after 12 days booster. Pie chart represent the fraction of the total CD4$^+$ IFNγ$^+$ cytokines response comprising any combination of TNF and IL-2 production after Flublok® stimulation in different immunized groups. Beneath pie chart, bar graph represents the frequencies of multifunctional T cells in CD4$^+$ IFNγ$^+$ T cell compartment. Statistical comparison employed test one-way ANOVA; *$p < 0.033$, **$p < 0.002$, ***$p < 0.001$ (n = 5 - 12 per group). Study was inclusive of two independent repeats.

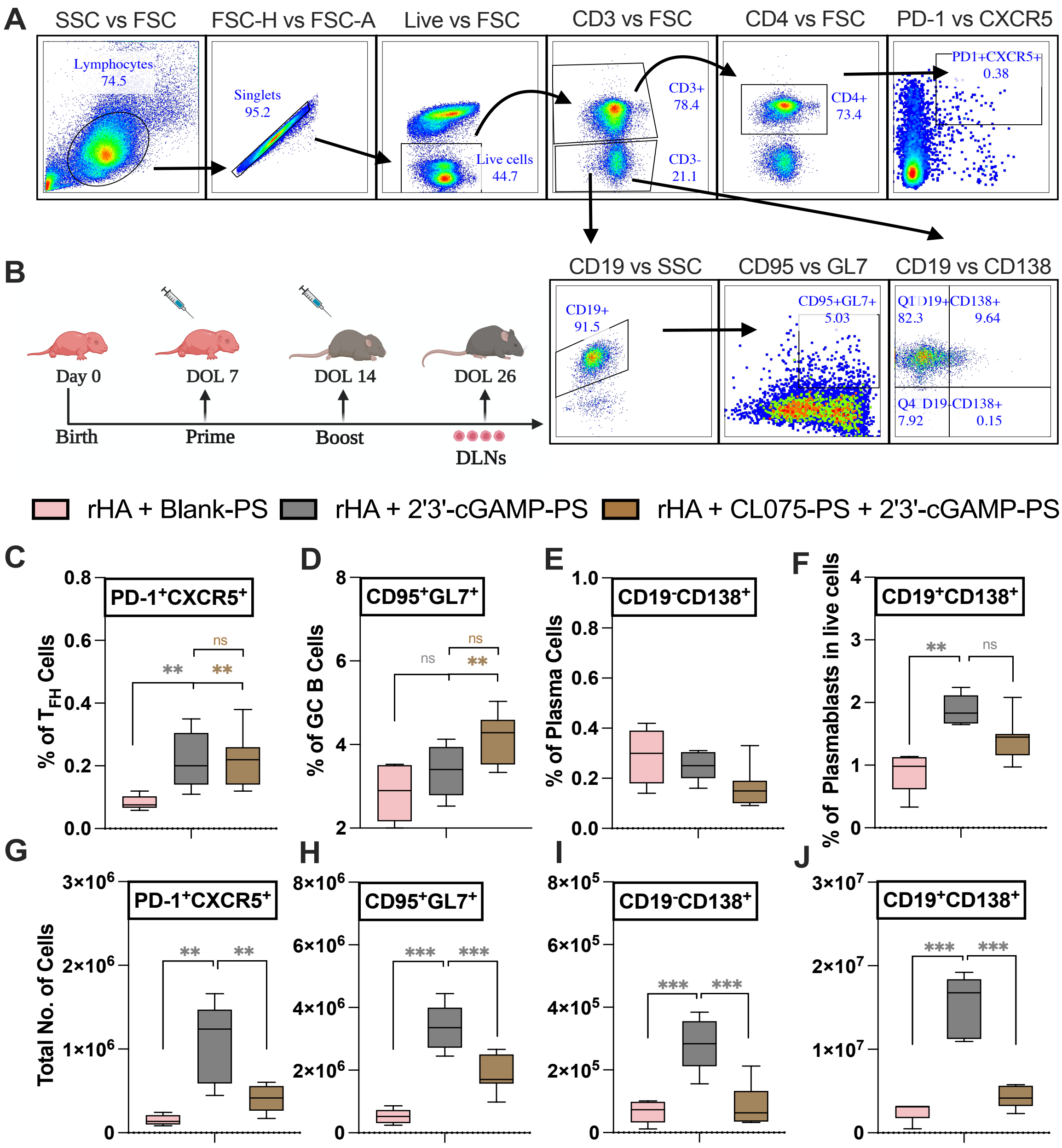


**Figure S4. cGAMP and CL075 encapsulating PS promotes $T_{FH}$ and B cell responses in draining lymph nodes (DLN).** (A) Shown was an example of the hierarchical gating strategy leading to the identification of $T_{FH}$ cells ($CD3^+CD4^+PD\text{-}1^+CXCR5^+$), GC B cells ($CD3^-CD19^+CD95^+GL7^+$), plasmablasts ($CD3^-CD19^+$ $CD138^+$) and plasma cells ($CD3^-CD19^-CD138^+$) are shown. (B) Infant C57BL/6 were immunized i.m. as described in Figure 4. Lymphocytes from DLN (both popliteal and inguinal) of immunized mice were harvested at DOL 26 for FACS analysis. (C) % of $T_{FH}$ cells, (D) % of GC B cells, (E) % of Plasma cells and (F) % of plasmablasts among live cells. (G-J) Total number of immune subsets in DLN. Statistical comparison employed test one-way ANOVA; $*p < 0.033$, $**p < 0.002$, $***p < 0.001$ (n = 5 - 7 per group), with comparison to rHA and mock loaded PEG-b-PPS nanocarriers control groups. Study was inclusive of two independent repeats.

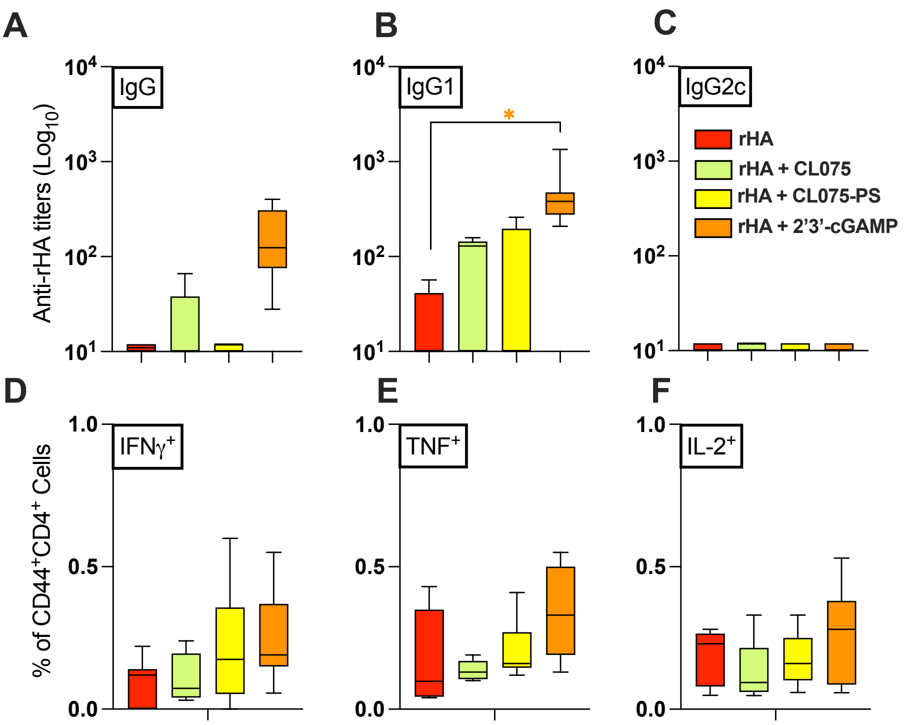


**Figure S5. Comparison of rHA specific humoral and cell mediated responses in between different formulations.** After immunization of infant C57BL/6 mice i.m. on DOL (day of life) 7 and 14, antibody titers for rHA-specific IgG (A), IgG1(B) and IgG2c (C) were determined by ELISA in serum samples collected at DOL 21. (D-F) Splenic $CD4^+$ T cell responses after rHA stimulation was determined by flow cytometry. Statistical comparison employed test one-way ANOVA; $^*p < 0.033$, $^{**}p < 0.002$, $^{***}p < 0.001$ ($n = 5 - 12$ per group). Study was inclusive of two independent repeats.

#### **Supplementary Table 1: Vaccine formulations for *in vivo* study**.

For the (rHA+Blank-PS) group, PS content was maintained at 400 μg to match the PS content of admixture group, i.e. rHA+CL075-PS+2’3’-cGAMP-PS.

| **Group** | **rHA (Flublok® Quadrivalent 2020-2021)** | **2’3’-cGAMP (μg)** | **CL075 (μM)** | **PS (μg)** |
|---|---|---|---|---|
| PBS | -- | -- | -- | -- |
| Blank-PS | 4μg (1μg of each variant) | -- | -- | 400 μg |
| CL075 | 4μg (1μg of each variant) | -- | 164 μM | -- |
| 2’3’-cGAMP | 4μg (1μg of each variant) | 1μg | -- | -- |
| CL075+2’3’-cGAMP | 4μg (1μg of each variant) | 1μg | 164 μM | -- |
| CL075-PS | 4μg (1μg of each variant) | -- | 164 μM | 200 μg |
| 2’3’-cGAMP-PS | 4μg (1μg of each variant) | 1μg | -- | 200 μg |
| CL075-PS + 2’3’-cGAMP-PS | 4μg (1μg of each variant) | 1μg | 164 μM | 400 μg |
| [CL075+2’3’-cGAMP]-PS | 4μg (1μg of each variant) | 1μg | 164 μM | 200 μg |

### Supplementary Table 2:

Panels and regents for flow cytometry assay.

| Assay | Target | Clone | Fluorochrome | Vendor | Identifier | Titer |
|---|---|---|---|---|---|---|
| | Viability | --- | eFlour780 | eBioscience | 65-0865 | 1:1000 |
| | CD3 | UCHT1 | PerCP-Cy5.5 | BioLegend | 300430 | 1:40 |
| T-Cell | CD4 | RPA-T4 | PE | BioLegend | 300508 | 1:40 |
| Cytokines | CD8 | RPA-T8 | PE-Dazzle 594 | BioLegend | 301057 | 1:40 |
| (Human) | IL-4 | MP4-25D2 | BV421 | BioLegend | 500826 | 1:40 |
| | IL-17A | N49-653 | Alexa Fluor 647 | BD Biosciences | 560491 | 1:10 |
| | IFN-$\gamma$ | B27 | Alexa Fluor 488 | BD Biosciences | 557718 | 1:40 |

| Assay | Target | Clone | Fluorochrome | Vendor | Identifier | Titer |
|---|---|---|---|---|---|---|
| | Viability | --- | Brilliant Violet 510 | Invitrogen | L34966 | 1:500 |
| | CD3 | 17A2 | Brilliant Violet 785 | BioLegend | 100232 | 1:40 |
| T-Cell | CD4 | RM4-5 | APC/Fire 750 | BioLegend | 100568 | 1:160 |
| Cytokines | CD8 | 53-6.7 | Brilliant UltraViolet 395 (BUV395) | BD Biosciences | 563786 | 1:80 |
| (Mice) | | | | | | |
| | CD44 | IM7 | PerCP-Cy5.5 | BioLegend | 103032 | 1:160 |
| | IFN-$\gamma$ | XMG1.2 | Alexa Fluor 488 | BioLegend | 505813 | 1:160 |
| | TNF | MP6-XT22 | PE Cy7 | BioLegend | 506324 | 1:160 |
| | IL-2 | JES6-5H4 | PE | BioLegend | 503808 | 1:40 |

| Assay | Target | Clone | Fluorochrome | Vendor | Identifier | Titer |
|---|---|---|---|---|---|---|
| | CD3 | 17A2 | BUV395 | BD Biosciences | 612803 | 1:80 |
| | PD-1 | 29F.1A12 | PE | BioLegend | 135206 | 1:40 |
| DLNs | CXCR5 | L138D7 | Brilliant Violet 421 | BioLegend | 145512 | 1:40 |
| immunophenotyping | CD19 | 1D3/CD19 | FITC | BioLegend | 152404 | 1:320 |
| (Mice) | CD95 | Jo2 | PE Cy7 | BD Biosciences | 557653 | 1:160 |
| | GL7 | GL7 | Alexa Fluor 647 | BioLegend | 144606 | 1:80 |
| | CD138 | 281-2 | Brilliant Violet 785 | BioLegend | 142534 | 1:80 |
